# Additively manufactured multiplexed electrochemical device (AMMED) for portable sample-to-answer detection

**DOI:** 10.1101/2023.08.17.553741

**Authors:** Arash Khorrami Jahromi, Roozbeh Siavash Moakhar, Sripadh Guptha Yedire, Hamed Shieh, Katia Rosenflanz, Amber Bricks, Justin de Vries, Yao Lu, Houda Shafique, Julia Strauss, Sara Mahshid

## Abstract

Portable sample-to-answer devices with applications in point-of-care settings have emerged to obviate the necessity of centralized laboratories for biomarker analysis. In this work, a smartphone-operated and additively manufactured multiplexed electrochemical device (AMMED) is presented for the portable detection of biomarkers in blood and saliva. AMMED is comprised of a customized portable potentiostat with a multiplexing feature, a 3D-printed sample collection cartridge to handle three samples of saliva and blood at the same time, a smartphone application to remotely control the potentiostat, and a 3D-printed-based multiplexed microfluidic electrochemical biosensor (test chip). Here, by employing additive manufacturing techniques, a simple, cleanroom-free, and scalable approach was proposed for the fabrication of the test chip. Moreover, these techniques can bring about easy integration of AMMED components. Additionally, the test chip can be compatible with different affinity-based bioassays which can be implemented in a multiplexed manner for detection. The AMMED components were successfully characterized in terms of electrochemical and fluidic performance. Particularly, to demonstrate the biosensing capabilities of the device, the spike protein of the SARS-CoV-2 omicron variant and a well-established aptameric assay were selected as the representative biomarker and the bioassay, respectively. The proposed device accurately and selectively detected the target of interest in a rapid (5 min) and multiplex manner with a dynamic detection range of 1–10,000 pg. ml^-1^ in different media; and the clinical feasibility was assessed by several saliva patient samples. AMMED offers a versatile sample-to-answer platform that can be used for the detection of various biomarkers present in biofluids.

## 1. Introduction

Human biofluids sampling (e.g., blood, urine, cerebral spinal fluid, sweat, or saliva) provides a well-established approach to differentiating healthy from diseased states by offering a broad range of specific measurable biomarkers.^1^ Blood, as a gold-standard source of biomarkers, and saliva, as a non-invasively accessible body fluid, are attractive clinical bio-media containing various biomarkers, which can be indicative of a wide range of health conditions.^2-9^ Interestingly, it has been reported that there are reasonable correlations between salivary and blood serum concentrations of numerous biomarkers, highlighting the feasibility of using saliva samples to obtain equivalent clinical information.^10^ Moreover, the simultaneous determination of single or multiple, but clinically relevant, biomarkers is of high importance in reliable, sensitive, and specific medical diagnosis.^11, 12^ As a result, rapid, accurate, and multiplexed detection of biomarkers using simple and equipment-free devices is of great significance in realizing an early-stage diagnosis and continuous health monitoring. To reach these goals, the challenges of multiplexed sample-to-answer platforms, integrating the whole process of detection from sample preparation, to signal amplifications, to readout displays in a point-of-care (POC) manner should be addressed.^13^ Sample-to-answer devices with potential applications in POC settings are mainly comprised of a (i) sample collection and handling system, (ii) quantitative assay, and (iii) transducer and readout.

Microfluidic techniques are commonly employed for precise fluidic control, low reagent consumption, and rapid reaction progression, enabling on-platform parallel sample collection and handling as well as multiplexed detection in a miniaturized form.^14, 15^ The micropumps which provide fluid displacement with minimum use of external power source/field or actuators (e.g., finger-powered,^16^ capillary-driven,^17^ and simple suction-based systems^18^) have superiority in terms of being compatible with POC devices.^19^ These kinds of microfluidics systems have also rapid and mass-producible manufacturing processes in most cases. Additive manufacturing (3D printing) techniques can substantially address the outstanding challenges associated with conventional microfabrication techniques (e.g., photoresist coating, photolithography, solvent-based lift-off) on a large scale.^20, 21^ Indeed, Additive manufacturing (AM) can bring about outcomes comparable to the traditional methods in terms of temporal and spatial resolution, material, and mechanical properties while being cost-effective, rapid, and scalable.

Label-free affinity-based quantitative assays has gained more attention compared to other biosensing approaches (such as biocatalytic assays) as this method offers the possibility of sensing a wide range of biomarkers with simplified design, lower reagent costs, and real-time measurements.^22, 23^ This method employ a non-reactive approach, in which specific binding interaction between the analyte and biorecognition elements (BREs) such as antibodies, aptamers, molecularly imprinted polymers (MIPs) is converted into a measurable signal by a transducer.^24, 25^ However, biosensing platforms implementing label-free affinity-based assays are encountered with two main challenges. First, these systems rely on a single binding event, resulting in the need to implement a BRE with extremely high affinity and specificity for the target. Secondly, the lack of significant signal amplification elements necessitates a highly sensitive transducer. In this regard, we can exploit the potentials of highly stable and selective BREs (such as aptamers) and nanostructured materials, and facilitating the electron transfer between the electroactive species and transducer. ^5, 6, 26^

Electrochemical transducers/readouts have garnered significant attention because of their relatively low cost, accuracy, low turnaround time, simplicity, and low instrumentation requirements, and importantly being appealing for multiplex POC biomarker detection.^27, 28^ However, traditional electrochemical biosensors remain limited to detection on peripheral equipment including standard reference and counter electrodes, as well as bulky benchtop potentiostats that cost >$10,000 and can only be operated by trained personnel. Portable variants of the benchtops (with similar specifications that costs still above 1000 USD) have been presented by several companies (e.g., Gamry, PalmSens, Ivium, and Metrohm). However, they provide limited information about their hardware and software technical specifications because they operate as black boxes, making the development of new measurement techniques and integration with other instruments more challenging.^29^ Several research groups developed homemade customized potentiostats with the potential use in POC. ^30-33^ However, the devices lack one or a few important capabilities to (i) run both voltammetry and EIS (ii) detect multiple analytes in a multiplexed manner (iii) be used for the determination of various concentrations of biomarkers in biological media, and real clinical samples. Also, the design of electrodes is another important aspect of reliable and multiplexed electrochemical biosensing, which can avoid cross-contamination and allow for the simultaneous detection of multiple analytes using either a single or several quantitative assays.

In the present work, a portable and smartphone-operated additively manufactured multiplexed electrochemical device (AMMED) with application in POC or point-of-use settings is proposed. AMMED was developed in a way to meet the POC testing criteria by employing a suction-based microfluidic system relying on a single-trigger mechanism alongside a filtered-based sample collection approach, a dimeric DNA aptamer-based gold nanostructured (GNS) electrochemical biosensor (GNS aptasensor), and a customized potentiostat. Indeed, the device mainly consists of a homemade smartphone-controlled portable potentiostat featuring the multiplexed option to run voltammetry and EIS measurements (AMMED potentiostat, **Fig. 1a(i)**), a smartphone application to remotely run the potentiostat and receive data (AMMED smartphone application, **Fig. 1a(ii)**), a kit to collect and handle saliva and blood samples (AMMED sample collection cartridge, **Fig. 1a(iii)**), with embedded paper-based filter to effectively reduces saliva viscosity that facilitates sample delivery, and a multiplexed microfluidic electrochemical biosensing platform employing the GNS aptasensor (AMMED test chip, **Fig. 1a(iv)**). Interestingly, the AMMED components were fabricated based on AM (3D-printing) techniques, making the whole process almost cleanroom-free and scalable. In particular, the test chip consisting of electrodes and microfluidic part was easily fabricated using a new method briefly described in **Fig. 1b** (see more details in **Fig. 2**). This approach obviates the necessity for time-consuming, challengeable, and expensive traditional methods such as photoresist coating, photolithography, and solvent-based lift-off. Furthermore, the main components of AMMED including the fluid sample handling system, the GNS aptasensor, and the customized homemade potentiostat were characterized and successfully evaluated in terms of their functionality. The biosensing performance of the proposed device was investigated by the detection of a diagnostic representative biomarker in the relevant biofluids, including saliva and diluted whole blood. Spike protein (S-protein) of the SARS-CoV-2 omicron variant was selected as the representative biomarker since it has been proven to be a valuable antigen for non-invasive, accurate, and rapid diagnosis of COVID-19.^34, 35^ Additionally, AMMED was challenged with several saliva patient samples.

**Fig. 1.**
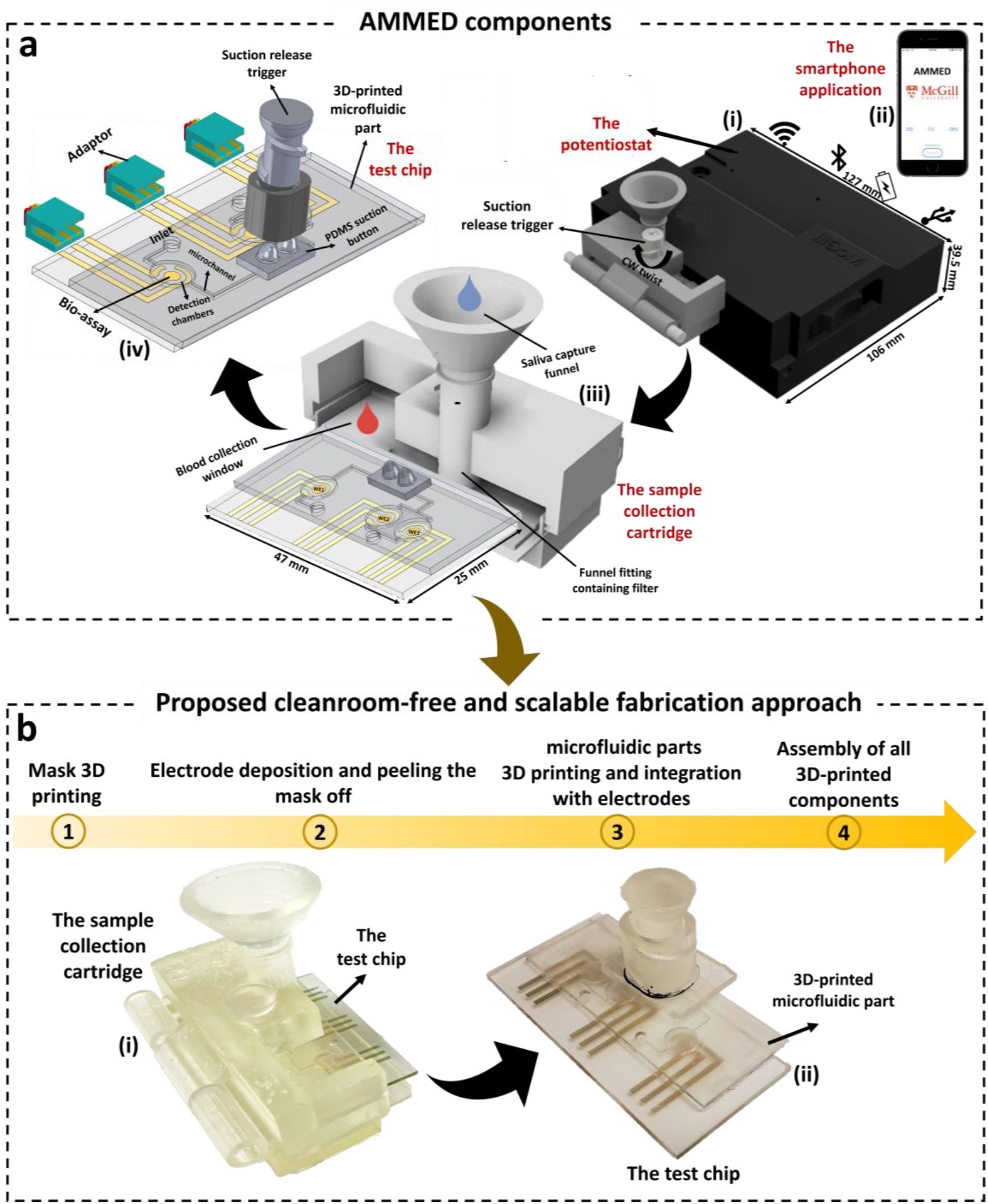
Representation of various components and fabrication process of AMMED. a) AMMED device scheme comprised of a (i) portable customized potentiostat, (ii) smartphone application, (iii) sample collection cartridge, (iv) test chip. b) Fabrication process flow of AMMED, based on additive manufacturing; 3D-printed-based (i) sample collection cartridge and (ii) test chip

**Fig. 2.**
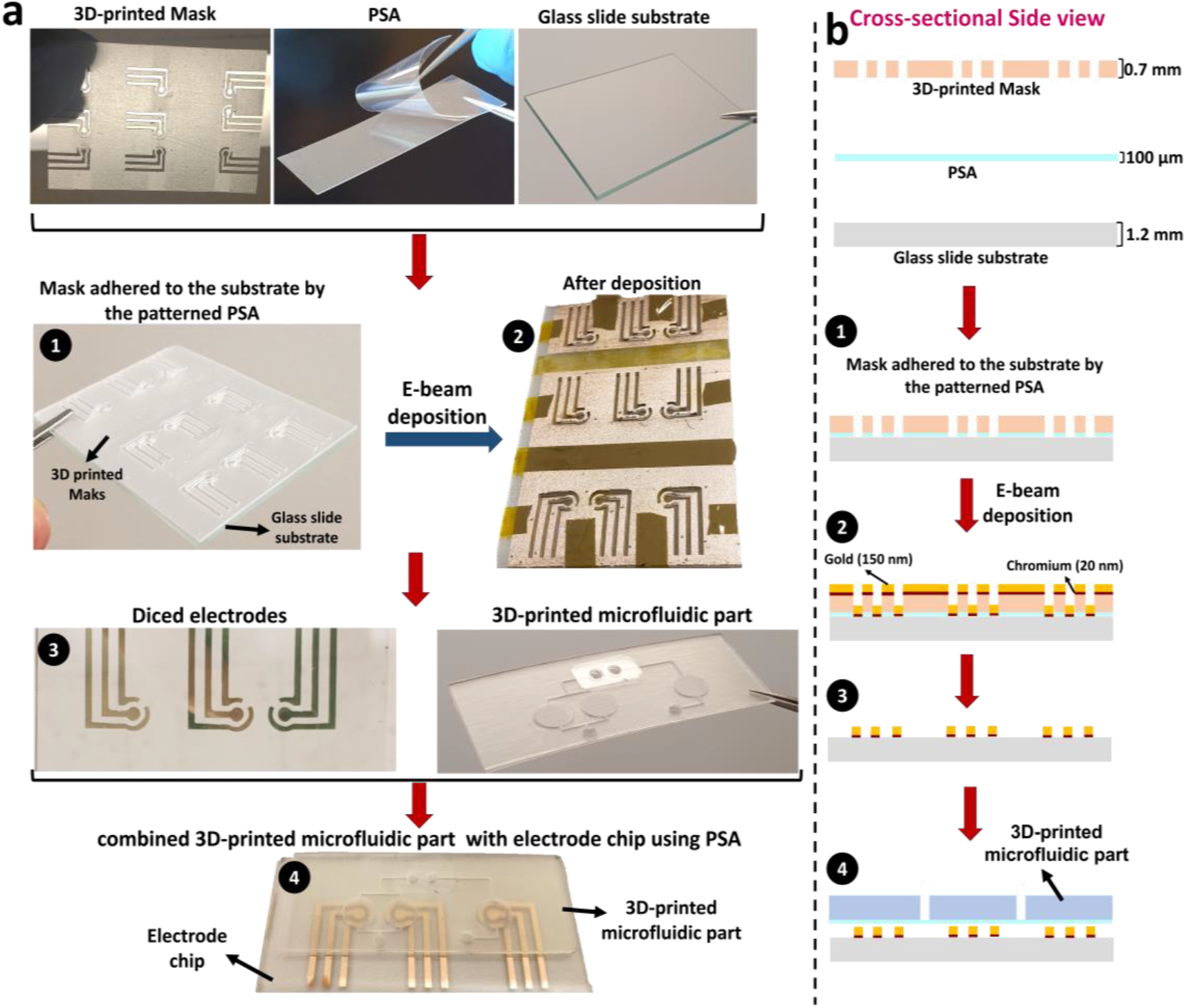
Proposed cleanroom-free and scalable fabrication process of AMMED test chip. a) Real images of each fabrication step. b) Schematic cross-sectional view of each fabrication step (b). 1) Fixing the 3D-printed mask featuring the electrode patterns onto the glass slide substrate using an adhesive. 2) Sequential deposition of chromium (adhesion layer), and gold onto the masked substrate for creating the electrodes. 3) Peeling off the 3D-printed mask and dicing the glass slide substrate into individual chips. 4) Mounting the 3D-printed microfluidic part onto the electrode chip using PSA for controlling sample delivery and limiting the sensing area.

## 2. Experimental Section

### 2.1. Reagents and materials

Gold (III) chloride trihydrate, Potassium hexacyanoferrate(III), Potassium hexacyanoferrate(II) trihydrate, Tris(2-carboxyethyl) phosphine hydrochloride (TCEP), 6-Mercapto-1-hexanol (MCH), and Phosphate buffer saline (PBS) 1X were purchased from Sigma-Aldrich Canada Co, Oakville, Ontario. All solutions were prepared using either ultrapure water (>18 MΩ cm) from a Millipore Milli-Q water purification system or PBS 1X. The viral proteins used in this work include SARS-CoV-2 B.1.1.529 (Omicron) spike S1 protein (40591-V08H41 SinoBiological), human coronavirus (HCoV-229E) spike protein (RDC3141; Cedarlane), Middle East respiratory syndrome coronavirus (MERS-CoV) spike protein (40069-V08B1; SinoBiological), Influenza A H1N1 HA protein (NBP1-99041; Cedarlane). Single-donor human whole blood and pooled human saliva were purchased from Innovative Research, Inc. The dimeric aptamers, proposed by Zhang, Zijie, et al.,^36^ with a sequence of 5’ – S1 - 20T – S2 – thiolC3 - 3’, where S1 = TTCCG GTTAA TTTAT GCTCT ACCCG TCCAC CTACC GGAA and S2 = ACGGG TTTGG CGTCG GGCCT GGCGG GGGGA TAGTG CGGT, with *K*_*d*_ = 0.17 ± 0.05 nM, were synthesized by Base Pair Biotechnologies and purified using HPLC technique. The screen-printed gold electrodes (SPGE, Dropsens, DRP 220AT) were supplied from Metrohm Canada, Mississauga, Ontario.

### 2.2. Fabrication of AMMED test chip and sample collection cartridge

**Fig. 2** shows the process for the fabrication of AMMED test chip based on an additive manufacturing scheme. **Fig. 2a** and **Fig. 2b** show real images of each fabrication step and schematic cross-sectional side view of the corresponding step, respectively. The test chip was fabricated based on a glass slide substrate. In the proposed fabrication scheme, a 3D-printed mask has been used for creating the desired pattern of electrodes. First, ethanol and DI water were used for cleaning the glass substrate prior to mask attachment and gold deposition. Then, a 700 μm thick 3D-printed mask (fabricated using SLA 3D-printer Form 3, Formlabs) featuring the patterns for the three-multiplexed electrodes was attached to the glass substrate using a pressure sensitive adhesive (PSA) film, resulting in **Fig. 2a1** and **Fig. 2b1**. This 3D-printed mask allowed for the fabless patterning of gold electrodes in a single step. Subsequently, a 20 nm film of Chromium, as the adhesion layer, was deposited onto the masked substrate followed by a 150 nm layer of gold using electron beam evaporator Temescal BJD 1800 (**Fig. 2a2** and **Fig. 2b2**). The 3D-printed mask was removed after the deposition and the glass slide substrate was diced into individual chips using a Disco DAD 3240 dicing saw (**Fig. 2a3 & Fig. 2b3**). Some optimizations were done in terms of the 3D-printed mask fabrication to obtain optimal mask and glass attachment, leading to more precise electrode deposition (see **section S1** and **Fig. S1**).

The microfluidic channels and detection chambers of the test chip were fabricated based on SLA 3D printing (at 50 μm resolution on the z axis). Finally, the 3D-printed microfluidic part was mounted onto the glass chip using PSA to control sample delivery, encapsulate the device, and confine the sensing area of electrodes (**Fig. 2a4** and **Fig. 2b4**). The suction button preparation and attachment to the test chip was described in **Fig. S2**. This was followed by GNS preparation on the deposited flat gold surface whose protocol was described in our previous works.^37, 38^ Once the GNS working electrodes (WEs) were electrodeposited, biochemical functionalization procedure involving aptamers immobilization and MCH modification was carried out to prepare the GNS aptasensor (see details in the following subsection).

The sample collection cartridge parts (**Fig. S3a**) were simply 3D printed using FormLabs resin (Form 3, Formlabs) with high resolution, followed by washing and curing steps (see **section S3** in SI). All designs were made on AutoCAD 2022 and Fusion 360 (Autodesk) or Inventor. For more details about the designs and layouts, readers are referred to open-source files accessible through **[GitHub link 1](https://github.com/justindevries/MEMA/tree/main)** (**section S8** in SI).

### 2.3. Biochemical functionalization of GNS WEs

Firstly, the lyophilized dimeric aptamers were reconstituted using Tris-EDTA (pH=7.5). Aptamer folding buffer (50 mM Tris-HCl buffer, 10 mM KCl, 100 mM NaCl, 50 mM MgCl2) was utilized to dilute the aptamer solution to 10x working concentration, and the solution was heated in an oven at 90-95°C for approximately 5 min. After cooling to room temperature (15 minutes), the thiolated aptamers were diluted by reducing buffer (1X PBS, containing 2 mM MgCl2, and TCEP 10 mM, 1:1 ratio) to 5x working concentration, and incubated for 120 min at room temperature. Subsequently, PBS, containing 1 mM MgCl2, was added to the reduced aptamer solution to make desired working concentration of the aptamer to be immobilized on the WE surface. To realize the optimal performance of the aptasensor, several concentrations of aptamer solution (0.5 μM, 1 μM, 2 μM, 4 μM, 5 μM) were tested. 15 μL of the aptamer’s solutions were incubated on the WE surface for an overnight in a 4°C fridge. Subsequently, 5 μL of MCH (100 mM) was used to cover the WE for 20 min to block non-functionalized areas, avoiding possible non-specific binding. Between each functionalization step, the surface of the WEs were washed using PBS (containing 1mM MgCl2), dried and electrochemically characterized.

### 2.4. AMMED potentiostat and smartphone application

AMMED potentiostat (**Fig. S3b**) makes use of a custom Printed Circuit Board (PCB) based largely on that of Jenkins et al.’s ABE-Stat, a recently developed EIS-compatible open-source potentiostat.^30^ Our modified design includes an updated Bluetooth protocol (Bluetooth Low Energy 5.1, called BLE) and the incorporation of a relay module to allow for multiplexing of multiple electrodes (see **section S3** and **Fig. S4**). The user interfacing is performed through a custom smartphone application, available for both Android and Apple devices (**Section S8, GitHub [links 2](https://github.com/juliamstrauss/NFluidEX-iOS)** and **[link 3](https://github.com/juliamstrauss/NFluidEX-Android)**) allowing for Electrochemical Impedance Spectroscopy (EIS), Cyclic Voltammetry (CV), and Differential Pulse Voltammetry (DPV) testing with options to customize input parameters. **Fig. S3c** shows the real image of the whole AMMED device.

### 2.5. Electrochemical Measurements

Different electroanalytical measurements, including CV, DPV, and EIS were run, in the presence of redox probe solution, via AMMED and PalmSens4 potentiostats. Different concentrations of the redox probe were prepared based on varying the total concentration of K_3_Fe(CN)_6_/ K_4_Fe(CN)_6_. For example, 5 mM redox probe solution is comprised of 2.5 mM of K_3_Fe(CN)_6_, 2.5 mM of K_4_Fe(CN)_6_, 0.1 M KCl, and 1 mM MgCl_2_ diluted in PBS 1x. The comparisons were conducted using well-established SPGEs. The measurement parameters and protocol have been mentioned in **section S4** in the SI.

### 2.6. Biosensing measurements

After preparing the GNS aptasensors, biosensing measurements were conducted using both AMMED and PalmSens4/gold screen printed electrode setup for the further comparison purposes. A serial dilution of protein solutions was performed by spiking PBS, saliva and 10x diluted whole blood media with S-protein to prepare concentrations of 0.1, 1, 10, 100, 1000, 10000 pg/ml. The aptasensors were tested with the redox probe after 5 min incubation time of 15 μL of each concentration. Importantly, after target solution incubation (which is 5 min), the electrode surface is washed, resulting in removing biofouling proteins possibly absorbed on the surface. Then, the DPV measurement is performed in the redox probe solution. In order to eliminate the possible noises and hence more clearly show the DPV trendlines associated with each S-protein concentration, 6-degree polynomial curves were fitted to data points obtained from AMMED for each concentration (**Fig. S7**).

## 3. Results and discussions

### 3.1. Operational principle of AMMED sample collection cartridge and test chip

The aim of this design is to encompass passively-driven techniques that do not require complex user interfacing. The test chip is fitted into the custom 3D-printed sample collection cartridge for POC sample collection, pre-treatment, and analysis. The design is separated into distinct regions for saliva and blood collection for multiplexed fluid sampling onto the inlets of the device (**Fig. 1a(iii)**). Although saliva is an optimal choice for diagnostic purposes, the matrix effects, caused by the presence of glycoproteins, can be a hurdle in microfluidic flow. ^39^ Conventional filter-based methods can result in fluid loss and although centrifugation is the optimal method for clinical studies, it is not amenable to the POC. As such, we propose the use of a paper filter for low fluidic loss with high filter efficiency that can effectively reduce saliva viscosity (determined by vibrating viscometer, SV-10) after treatment using whatman grade 4 filter (**Fig. 3d(i), Fig. 3d(ii)**). This filter can be embedded within the sample collection cartridge for on-platform biofluid processing. POC fluid displacement was achieved through suction-based microfluidics using flexible PDMS buttons made from 3D printed molds in a soft lithography protocol. A single-trigger release enabled flow in both saliva and blood channels by tuning the dimensions of the suction buttons according to the volume of fluid needed to be displaced. Upon compression of the single trigger release, the suction buttons are compressed while fluid was loaded at the inlet (**Fig. 3a(i-iii)**). Following a single clockwise twist of the trigger, the compression of the flexible buttons are released to displace the fluid in a method resembling negative pressure allied to an eyedropper (**Fig. 3b(i-iii)**), thereby enabling unilateral flow across the microchannels of the test chip as shown in **Fig. 3c. Fig. 3C** demonstrates fluid motion (in form of food dye), driven by the suction-based micropump, where filled microchannels and the color distinction show no leaking or mixing between the samples which is an important factor for multiplexed detection. **Movie. S1** demonstrates how three liquid samples can be simply transferred from the inlet into the detection area (where GNS s WEs have been modified with the specific aptamers) by applying a suction trigger.

**Fig. 3.**
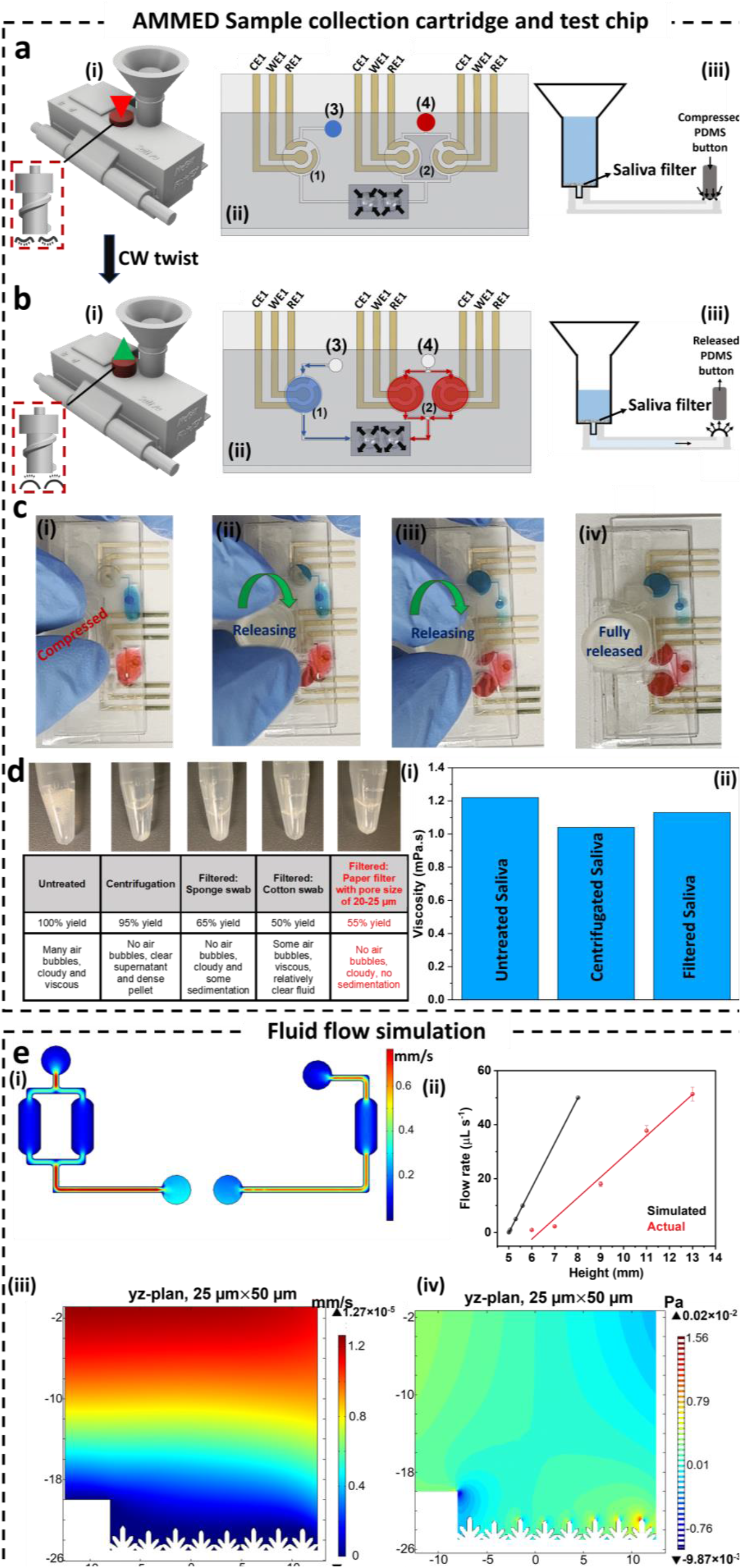
AMMED sample collection cartridge, test chip, and fluidic simulation. a) Initial position of the sample collection cartridge and the test chip; (i) schematic representation of the suction release trigger position, (ii) top view of the test chip before sample suction, 1) detection chamber for blood sample, 2) sensing area for saliva sample, 3) blood sample inlet (blood collection window), 4) saliva sample inlet (outlet of the funnel), (iii) cross sectional view of the saliva capture funnel with compressed PDMS suction button (initial position: no flow). b) Second position of the sample collection cartridge and the test chip; (i) schematic representation of the suction release trigger position, (ii) top view of the test chip after sample suction, (iii) cross-sectional view of the saliva capture funnel with released PDMS suction button (liquid suction). c) Food dye demonstration of fluid motion driven by on-chip micropump. d) (i) Comparison between different saliva treatment methods and filter selection, (ii) viscosity values for untreated, centrifugated and filtered (selected) saliva samples. e) Fluid flow simulation; (i) finite element analysis of the suction-based flow in the test chip showing 2D microchannels with the xy-plane velocity distribution (axis shown in millimeters; a 50 μm by 25 μm cross-sectional 2D profile of the yz-plane), (ii) actual and simulated flow rate in various saliva height in the column of the saliva capture funnel, (iii) velocity distribution and, (iv) pressure distribution in the chambers containing the aptamer-based electrochemical assay.

The suction-based flow was characterized with a finite element method simulation in COMSOL Multiphysics by assuming negative pressure applied over the microchannels (see **section S4**). The 2D flow profiles were evaluated assuming the creeping flow of incompressible fluids, showing the xy-plane velocity profiles (**Fig. 3e(i)**), that serve to demonstrate the low velocity in the detection chamber, minimizing the risk of detachment of the immobilized aptamers. Cross-sectional yz-plane profiles were evaluated under suction-based flow to demonstrate that both the incident velocity (**Fig. 3e(iii)**) and pressure (**Fig. 3e(iv)**) distributions remain low near the GNS s structures where detection is performed. The automated sample flow over the saliva collection funnel can also be simulated and characterized experimentally. Briefly, the sample was collected directly on the cartridge; the pressure drop to displace fluids across the filter and funnel was approximated using Darcy’s Law and a sudden contraction by considering the mechanical energy lost through friction (see **section S4** in SI). For the desired flow rate of 100 μL s^-1^ and known values for saliva viscosity, filter thickness, and filter permeability, we found a required pressure drop that can be modulated according to the height of fluid collected in the funnel, yielding the minimum required height of the designed self-collection funnel for passively driven flow to be 11 mm (**Fig. 3e(ii)**). It necessitates a minimum saliva volume of ∼ 700 μL, which is below the required volume by other saliva-based diagnostic collection protocols ^40, 41^, thereby demonstrating the advantage of a miniaturized device with on-platform biofluid collection.

### 3.2. AMMED potentiostat and smartphone application

AMMED potentiostat is compatible with SPEs via three evenly spaced SPE adaptors fitted into the front of AMMED test chip (**Fig. 4a**). From the frontward orientation depicted in **Fig. 4a**, this setup was designed specifically with our custom test chip in mind for blood-saliva testing, with the capability to run multiplexed electrochemical measurements on three sets of on-chip electrodes (WE, RE, and CE) at the same time. **Fig. 4b-c** illustrate the interior assembly of the potentiostat, which comprises a main PCB with an accessory relay module for simultaneous multiplexing and a manual single pole double throw (SPDT) switch for selection between single and multiplexed measurements. The PCB has three main states: calibration mode, ready for measurement, and measurement in progress, as indicated by the color of the on-board status LED (**Fig. 4d**). When in calibration mode, various resistors of known resistance values can be connected between the PCB’s WE and RE terminals for calibration of various sensing parameters, such as the transfer function of the digital-to-analog converter (DAC), the gains of the transimpedance amplifier (TIA), and the settings of the network analyzer. The precise calibration protocols can be found in Jenkins et al.’s supporting calibration guide.^30^ In ready for measurement state, electroanalytical instructions can be sent to the potentiostat over Bluetooth via our custom smartphone application, available for both Apple and Android devices (**Fig. 4e**, see **GitHub [links 2](https://github.com/juliamstrauss/NFluidEX-iOS) and [link 3](https://github.com/juliamstrauss/NFluidEX-Android)**). After starting up the app and connecting to AMMED potentiostat, the type of electrochemical technique can be selected (EIS, CV, or DPV), and measurement parameters (such as peak amplitude, start and end voltages, and scan rate) can be set. Once the message is received by the PCB, the potentiostat begins executing the communicated instructions, as indicated by the red status LED, and the measured current responses are transmitted back to the app in real-time. Once complete, a plot of the recorded data for the corresponding electrochemical technique is output to the user interface (**Fig. 4f**). The application uses Bluetooth low energy for transmitting testing instructions and receiving the resulting data from the device, allowing the platform to be used wirelessly and without WiFi. To integrate multiplexing capabilities, the application allows the user to choose between one or two, or three samples and analyzes the electrochemical data accordingly. A summarized flowchart for the app’s operation is shown in **Fig. 4g** (see **Fig. S3** for more details).

**Fig. 4.**
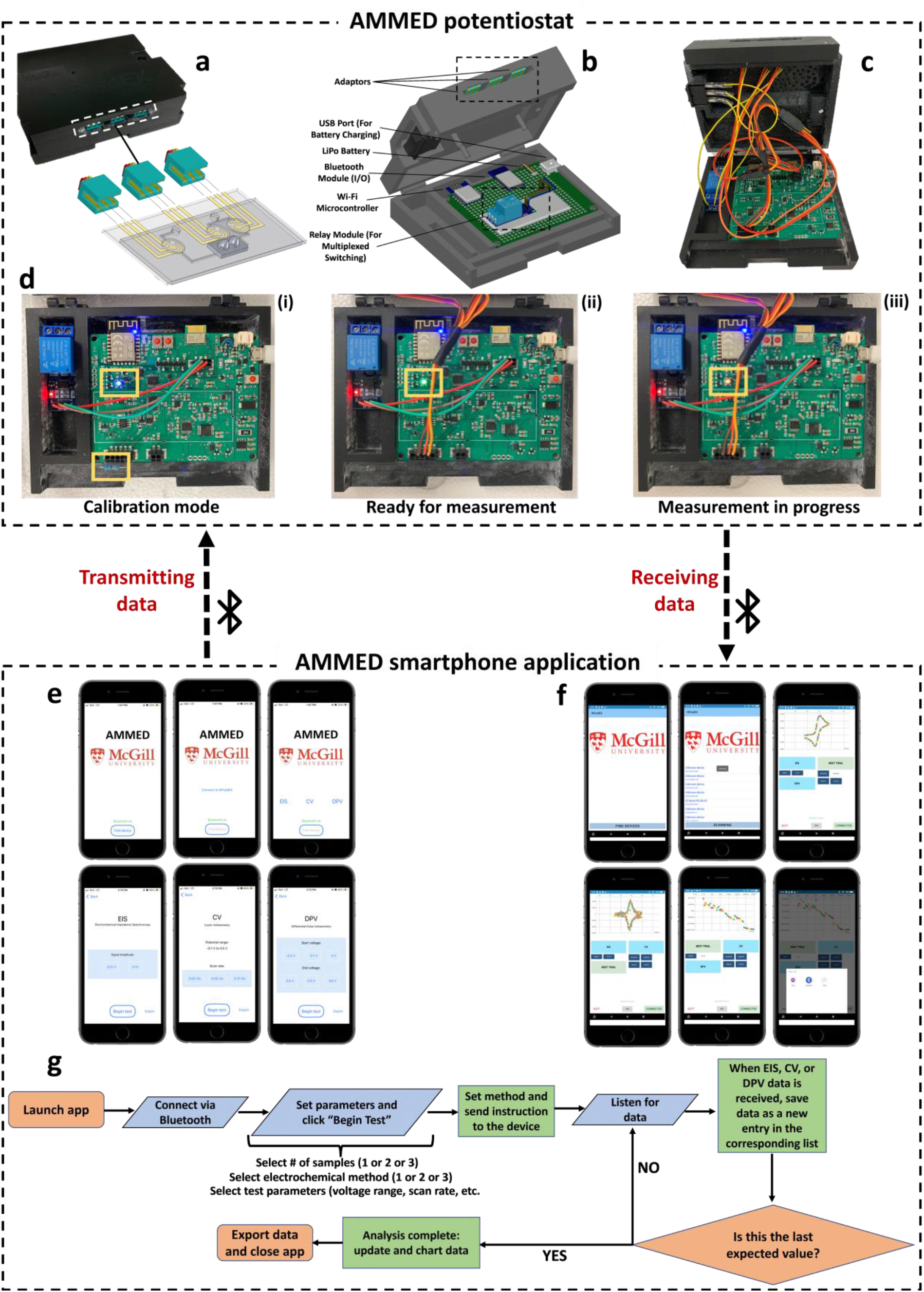
AMMED potentiostat and smartphone application. a) Configuration of the potentiostat connection to the test chip. b) Labeled diagram of AMMED potentiostat with the PBC and relay module. c) Interior view of AMMED potentiostat fully assembled with all device components. d) Operational states of AMMED potentiostat PCB; (i) calibration mode, (ii) ready for measurement, (iii) measurement in progress. e) Setting electroanalytical techniques and relevant parameters (transmitting data). f) Receiving results data. g) Summarized flowchart associated with the algorithm developed for the AMMED smartphone application.

**Table S1** comprehensively compares AMMED (a modified version of ABE-Stat) with existing portable potentiostat devices in terms of affordability and technologies. In addition to the superiority of the developed potentiostat mentioned in Table S1, this device has the feature to be easily integrated with the sample collection cartridge and test chip for multiplexed biosensing applications. Furthermore, the cost breakdown for the whole AMMED device has been summarized in **Table S3**. The performance of AMMED potentiostat (in terms of its comparison with commercial ones and sensing capability) was not disclosed in our previous works. Indeed, in our previous work, just a conceptual design of the potentiostat was suggested. Here, the potentiostat was successfully characterized (CV, DPV, and EIS) and its biosensing potential was demonstrated **(Figs. 4, Fig. 5**, and **Fig. 6**)

**Fig. 5.**
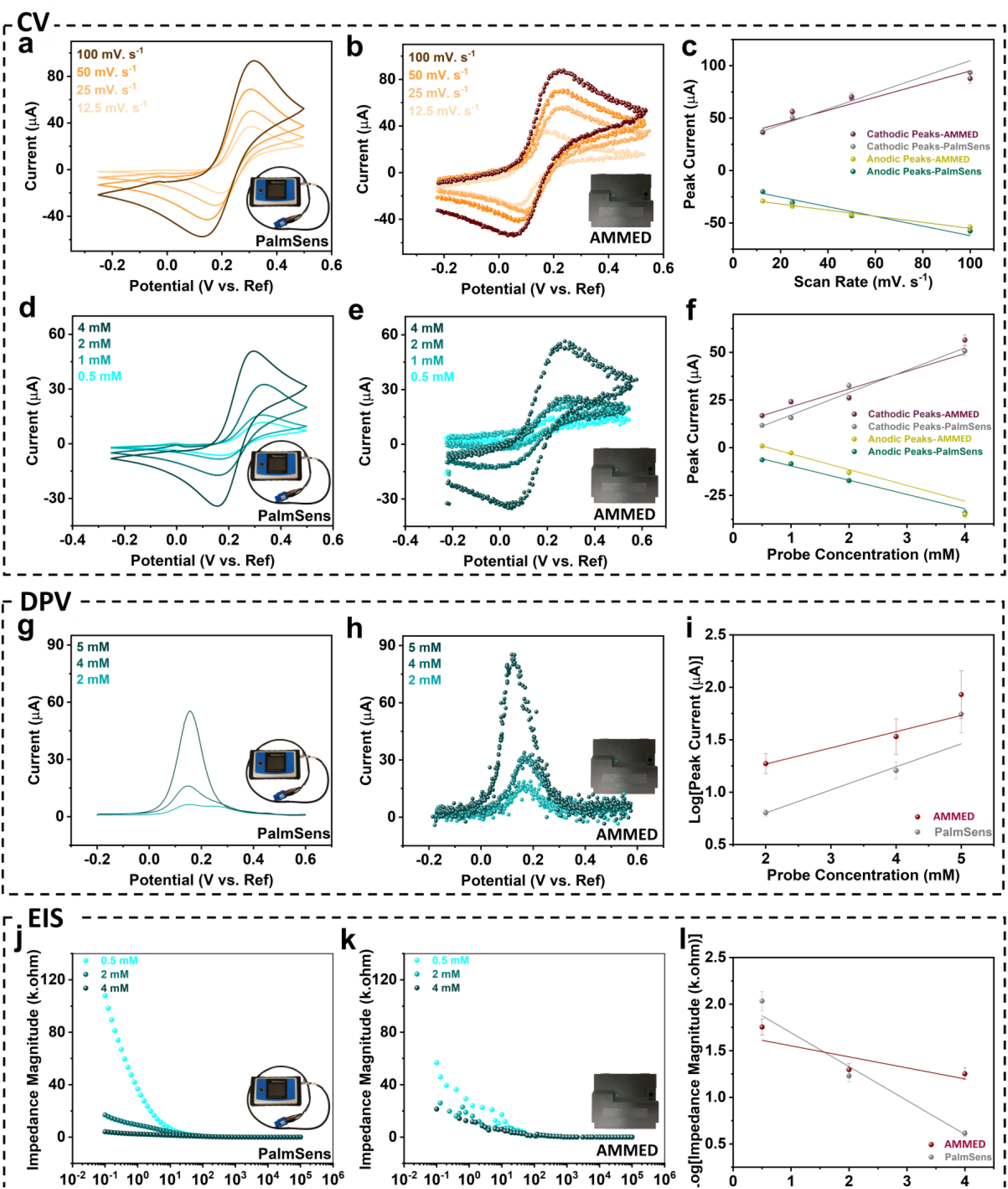
Electrochemical performance comparison between commercial PalmSens and AMMED potentiostat. The CV of the redox probe (5mM) at different scan rates recorded using (a) PalmSens and (b) AMMED; (c) the correlated linear relationship between the peak currents and the scan rates. The CV of different redox probe concentrations (at scan rate of 50 mV. S^-1^) recorded using (d) PalmSens and (e) AMMED; (f) the correlated linear relationship between the peak currents and the probe concentration. The DPV of different redox probe concentrations recorded using (g) PalmSens and (h) AMMED; (i) the correlated linear relationship between the peak currents and the probe concentrations. The impedance magnitude readout of different redox probe concentrations recorded using (j) PalmSens and (k) AMMED; (l) the correlated linear relationship between the impedance magnitudes at 10 Hz and the probe concentrations.

**Fig. 6.**
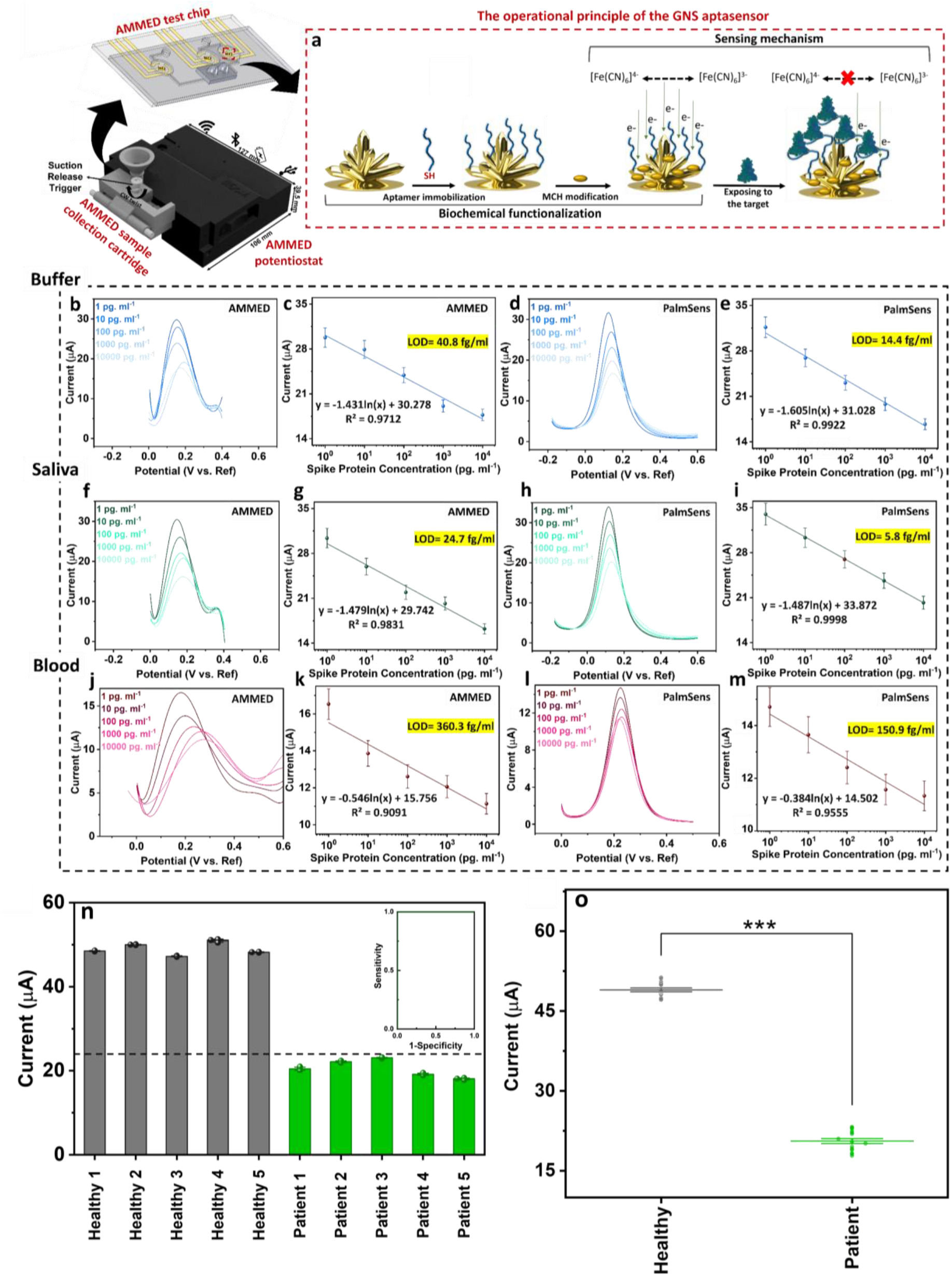
Analytical performance of AMMED and comparison with commercial PalmSens potentiostat, and Assessment of AMMED biosensing platform for the detection of SARS-CoV-2 in saliva samples using COVID-19-positive and -negative subjects. a) Schematic representation of the GNS aptasensor preparation, including the aptamer immobilization and MCH modification, followed by the aptaensor exposure to SARS-CoV-2 S-protein for biosensing evaluation. The DPV readout of SARS-CoV-2 S-protein on GNS aptasensors in different concentrations in buffer obtained from (b) AMMED and (d) PalmSens and (c) and (e) their corresponding calibration plots and LOD, respectively. The DPV readout of SARS-CoV-2 SP on GNS aptasensors in different concentrations in saliva obtained from (f) AMMED and (h) PalmSens and (g) and (i) their corresponding calibration plots, respectively. The DPV readout of SARS-CoV-2 SP on GNS aptasensors in different concentrations in blood obtained from (j) AMMED and (l) PalmSens and (k) and (m) their corresponding calibration plots, respectively. Data shows mean values ± standard deviation (n=3). (n) The DPV signal analysis of saliva samples comparing healthy and patient signals, with inset the ROC curve showing 100% sensing efficiency and (o) null comparison demonstrating the distinguished signal level in patients with COVID-19-positive compared to healthy controls.

### 3.3 Characterization and analytical performance of AMMED

#### 3.3.1 Electrochemical performance of AMMED potentiostat

The performance of AMMED in generating electrochemical signals was compared with signals obtained from commercial PalmSens 4 potentiostat. The comparisons were made via a series of electrochemical measurements including CV, DPV, and EIS. It is worth noting that no data smoothing was done to present the data obtained from AMMED. We conducted CV using the redox probe at four different scan rates including 12.5 mVs^-1^, 25 mVs^-1^, 50 mVs^-1^, and 100 mVs^-1^. **Fig. 5a**-**b** show the voltammograms recorded by PalmSens and AMMED, respectively. Regarding peak potential and current, AMMED plots resemble those from PalmSens. As can be seen in **Fig. 5c**, peak currents show an upward linear trend respect to scan rate, confirming diffusion-controlled behavior. Further, we studied the electrochemical performance of AMMED in the presence of the redox probe at different probe concentrations using CV and DPV. **Fig. 5d** and **5g** as well as **Fig. 5e** and **5h** demonstrate the corresponding CV and DPV voltammograms recorded by PalmSens and AMMED, respectively. As expected, CV and DPV show an increase in redox oxidation peak currents by increasing redox probe concentrations (**Fig. 5f** and **5i**). In a similar manner, increasing redox probe concentrations decrease redox reduction peak current. **Fig. 5j-k** show the Bode plots recoded by PalmSens and AMMED in the presence of different concentrations of redox molecules from 0.5 mM to 4 mM, respectively. AMMED recorded similar peak currents, potentials, peak shapes and impedance to the commercial device. As the commercial device is a black box, it is unknown whether signal filtration or processing occurs after data is collected. As previously mentioned, AMMED datasets did not undergo smoothing. They differ slightly, most likely due to the variations between electrodes and different PCB designs.

#### 3.3.2 Operational principle and characterizations of the GNS aptasensor

The GNS aptasensors, employed in AMMED test chip, were developed by combining a dimeric DNA aptamer-based assay with GNS s WE (described in our previous works^5^) in an electrochemical format. Each step of the biochemical functionalization process was electrochemically verified using DPV and EIS methods (**Fig. S5a**). As it was expected, decreased DPV peaks (**Fig. S5b**) and increased semicircle domain of the Nyquist plots (**Fig. S5c)** showed that both aptamer immobilization and MCH modification led to increased charge transfer resistance due to partially blocking electron transfer from the redox species. After preparing the GNS aptasensor, it was exposed to various concentrations of SARS-CoV-2 S-protein (0.1–10,000 pg. ml^-1^) spiked in PBS to assess its biosensing performance. EIS (**Fig. S5d**) and DPV (**Fig. S5f**) generated consistent differentiable signal changes. The increase in S-protein concentration resulted in higher charge transfer resistance, extracted from the Nyquist plot (linearly correlated in **Fig. S5e**), and similarly lower peak currents of DPV responses (linearly correlated in **Fig. S5g**). A hindrance effect caused by the aptamer/ S-protein complex disrupts the normal flow rate of redox probe toward the electrode surface of the GNS aptasensor, resulting in a decrease in electrochemical response and interfacial electron transfer. A hindrance effect may be caused by two factors: 1) S-protein binding to aptamers may restrain diffusion of the redox molecule; 2) the conformational changes associated with the folded aptamers upon the target capturing can also result in an additional steric hindrance. As a result, the higher the concentration of S-protein, the greater the steric hindrance effect is observed. In this step, all the measurements were performed by PalmSens4. The selectivity of the GNS aptasensor was evaluated in the presence of similar viral surface proteins, including HCoV-229E spike protein, MERS-CoV spike protein, and Influenza A H1N1 HA protein. The electrochemical response was conferred by the peak currents of the DPV plots and normalized to a relative signal, which yielded statistically significant (*p* < .001) differences when the aptasensor was incubated with the target of interest versus the interferant proteins in buffer (**Fig. S8a**), saliva (**Fig. S8b**) and blood (**Fig. S8c**).

#### 3.3.3 Multiplexed detection of SARS-CoV-2 S-protein using AMMED

Once the excellent performance of AMMED test chip and potentiostat were verified, we further evaluated the multiplexed detection performance of AMMED through biosensing the SARS-CoV-2 S-protein in relevant bio-media. AMMED DPV readouts were observed comparable with the results obtained from the combination of the GNS aptasensors and PalmSens potentiostat, as a benchmark. The integrated device consisting of AMMED test chip and potentiostat featuring multiplexed detection as well as the benchmark were tested to determine SARS-CoV-2 S-protein ranging from 1–10,000 pg. ml^-1^ in buffer (**Fig. 6b-e**), saliva (**Fig. 6f-i**) and blood (**Fig. 6j-m**). The decrease in the current signals were positively correlated to the increase in the concentration of the target analyte via PalmSens (**Fig. 6d-e, 6h-i, 6l-m**) and AMMED (**Fig. 6b-c, 6f-g, 6j-k**). As described in the experimental section, data smoothing was performed on AMMED data points for each concentration to more clearly show the DPV trendlines (**Fig. S9**). The signal difference remained negligible and within the same order of magnitude. As shown by the calibration curves, the linear detection range of the aptasensor recorded by the AMMED potentiostat is determined to be 1 pg/mL to 10,000 pg/ mL for detecting SARS-CoV-2 S-protein in buffer, saliva and blood media with the limits of detection below 400 fg/mL (see **section S6** in SI), confirming the reliable response of AMMED device in complex media as well as its capacity for yielding ultra-low LODs comparable with commercial readouts.

The results presented in **Fig. S7 f-g** were obtained using the commercial PalmSens 4, and they demonstrate the sensitivity and capability of our developed assay in detecting very low concentrations of SARS-CoV-2 S-proteins, as low as 0.1 pg. ml^-1^. These findings highlight the potential clinical relevance of our assay for early and accurate detection of the SARS-CoV-2 virus. However, in **Fig. 6**, our aim was to compare the results obtained from the AMMED with those obtained from the commercial PalmSens 4, showing that both devices yield comparable and reliable results. This comparative analysis validates the robustness and versatility of our assay and AMMED test chip and potentiostat, offering a promising approach for efficient and precise SARS-CoV-2 protein detection in various settings.

#### 3.3.4 Evaluation of AMMED performance using real clinical samples

AMMED device was further challenged with real saliva samples belonging to patients who had previously tested positive (n=5) or negative (n=5) on the standard polymerase chain reaction (PCR) diagnostic tests (**Fig. 6n**). Current responses were recorded from AMMED potentiostat on the GNS /aptasensor assay and yielded lower currents in all the COVID-19-positive samples compared to their COVID-19-negative counterparts. The difference in current was statistically different (**Fig. 6o**, *p* < .001) between patient and healthy samples on this portable multiplexed alternative with 100% sensitivity and specificity over the population of saliva samples (inset of **Fig. 6n**). It is common to report clinical responses via electrochemical signal readouts, and many other clinical and translational studies also do so. The reported currents in the calibration plot correspond to spike samples with varying SARS-CoV-2 S-protein concentrations. After MCH treatment, control samples displayed higher currents (∼40 microA). Healthy samples without SARS-CoV-2 S-proteins showed a similar response in the range of 47-51 microA. Readers are referred to section **Sections S6** and **S7** for more details about statistical analysis and COVID-19 patient samples.

## 4. Conclusion

The demand for POC biomarkers quantification continues to grow with the purposes of early-stage diagnosis, personalized medicine, and health monitoring. In the present work, the technological advancements for the realization of an easily manufactured and sample-to-answer device, named AMMED, for multiplexed detection of biomarkers in blood and saliva were successfully explored. A novel combination of a homemade portable potentiostat remotely controlled by an accompanying smartphone application, an additively manufactured filtered-based single-trigger sample collection cartridge, and a test chip, consisting of three microfluidic-based electrochemical GNS aptasensors, forms the AMMED device. Importantly, All the AMMED components, particularly the test chip which requires complicated microfabrication techniques based on traditional methods, were fabricated in a cleanroom-free and scalable way by implementing simple 3D-printing approach. The EIS and voltammetry measurements performed by AMMED potentiostat were validated with PalmSense4, as a commercial portable potentiostat. We demonstrated the clinical application of AMMED using a case study for the diagnosis of SARS-CoV-2 S-protein, as an important diagnostic biomarker for COVID-19. Various concentrations of S-protein (from 1pg/ml-10000 pg/ml) in buffer, saliva, and diluted whole blood media were determined in a multiplexed manner using AMMED with the limits of detection less than 90 fg. ml^-1^. The sensing performance of AMMED was comparable with the results obtained from a well-established electrochemical biosensing setup. Moreover, the selectivity studies showed that the proposed device was able to differentiate SARS-CoV-2 S-protein in different media from other related control analytes. Importantly, we realized successful clinical validation using several negative and positive real saliva samples of COVID-19. AMMED is the first of its kind to offer a versatile sample-to-answer platform that can be used for various biomarkers present in blood and saliva once its aptamer-based assay is modified according to the targets of interest.

## Supporting information

https://mcgill-my.sharepoint.com/:w:/r/personal/arash_khorramijahromi_mcgill_ca/Documents/Device/Manuscript_new%20-%20Preprint.docx?d=w9d6298358824495

## Acknowledgments

The authors thank the Faculty of Engineering at McGill University, the Natural Science and Engineering Research Council of Canada (NSERC, G247765), the Canadian Institutes of Health Research (CIHR, 257352), the Canada Foundation for Innovation (CFI, G248924), and MI4 Emergency COVID-19 Research Funding (250611), for financial support. The authors acknowledge Nanotools-Microfab and the Facility for Electron Microscopy Research at McGill University and the research facilities of NanoQAM at the Université du Québec à Montréal. CdRM thanks CONACYT (924317) and McGill Engineering Doctoral Award for the scholarship funds. This publication was supported by University Health Network’s PRESERVE-Pandemic Response Biobank for coronavirus samples, UHN Biospecimen Services, REB # 20-5364. Its contents are solely the responsibility of the authors and do not necessarily represent the official views of the University Health Network.

