## Supplementary material for "Additively manufactured multiplexed electrochemical device (AMMED) for portable sample-to-answer detection": https://mcgill-my.sharepoint.com/:w:/r/personal/arash_khorramijahromi_mcgill_ca/Documents/Device/Manuscript_new%20-%20Preprint.docx?d=w9d6298358824495

#### S1. Optimization of electrodes fabrication

We tested both flexible (flexible 80A, v1, Formlabs) and regular clear resin (clear resin v4, Formlabs) to fabricate the mask. We observed better patterning of the electrodes with the clear resin. Then, we investigated the optimum thickness of the mask, resulting in better attachment (without bending issues) to the glass substrate and hence acceptable electrode patterning. Based on our observation the mask with 0.7 mm thickness led to a more precise electrode patterning with higher repeatability and reproducibility (**Fig. S1**).

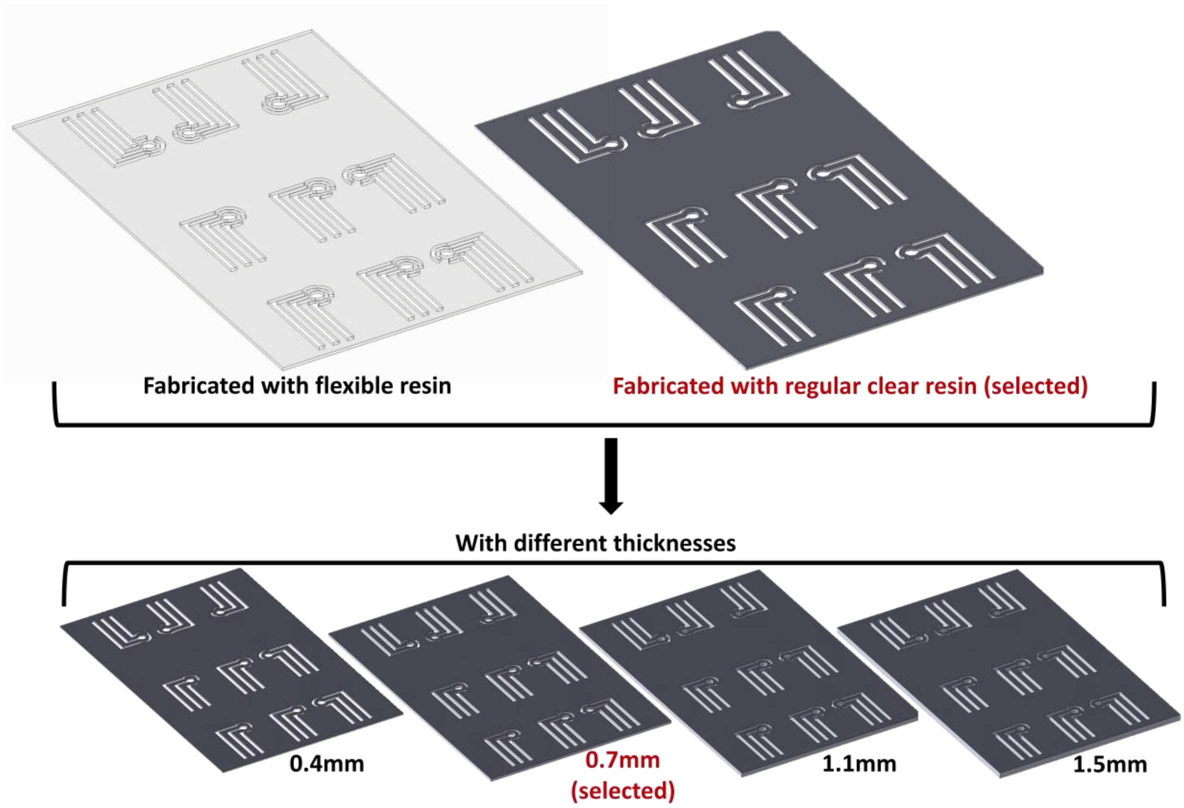


**Fig. S1. 3D-printed mask optimization for electrodes fabrication.**

### S2. Fabrication of AMMED sample collection cartridge

The sample collection cartridge parts (**Fig. S3a**) were simply 3D printed using FormLabs resin (Form 3, Formlabs) with high resolution, followed by placing the printed parts into FormLabs washer (where they are sonicated for 20 minutes in isopropyl alcohol to remove excess uncured resin) and FormLabs UV curing machine for 15 minutes at 60°C. All designs were made on AutoCAD 2022 and Fusion 360 (Autodesk) or Inventor. The sample collection cartridge features a saliva self-collection funnel lined with a Whatman Grade 4 filter (pore size: 20-25 µm) for sample pre-treatment to reduce biofluid matrix effects. The test chip is placed at the base of the sample collection cartridge and is firmly closed by rotating the hinges onto the chip with secured guide rails; the base of the funnel features a sudden contraction that fits directly into the microfluidic inlet. Blood collection follows the same protocol as a glucometer via a finger prick; therefore, a blood collection window exposes the microfluidic inlet for direct patient blood droplet collection. Regarding the elastomeric chamber fabrication protocol (**Fig. S2**), the elastomeric chambers are fabricated using SLA-printed molds. The 3D-printed molds are first surface treated to avoid curing inhibition at the PDMS mold interface. The high resolution of the printing ensured low roughness of the PDMS thereby ensuring strong bonding.

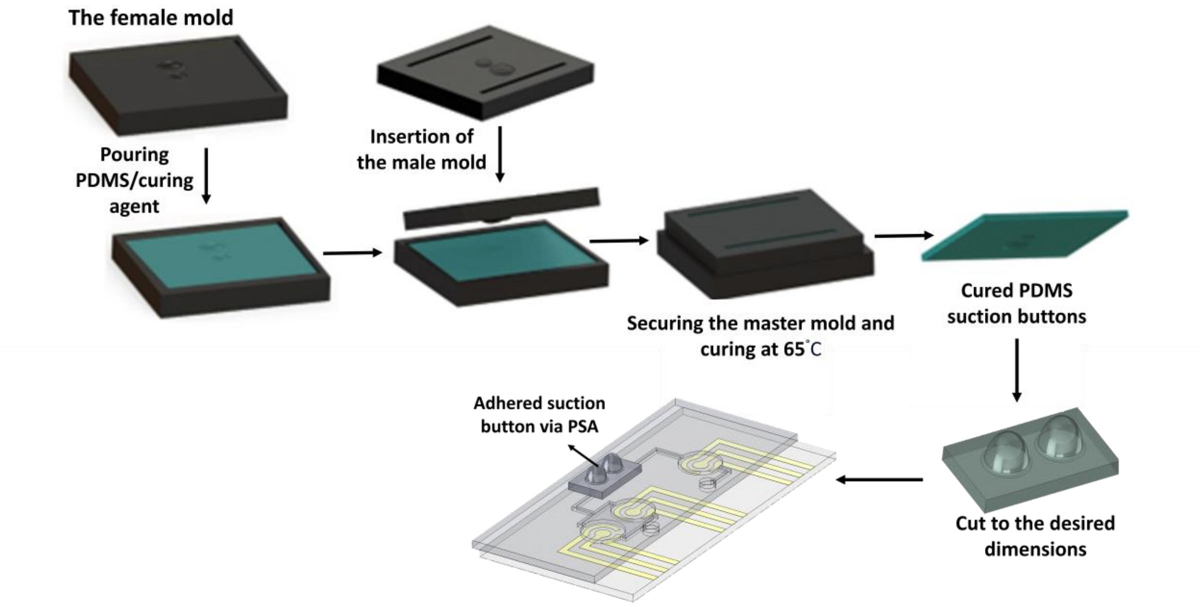


**Fig. S2. The suction button preparation procedure.**

**S3. AMMED potentiostat and smartphone application**

The device is controlled via an ESP-12F WiFi microcontroller (Espressif Systems, Shanghai, China) which receives user-input signals from a CYBLE-014008-00 Bluetooth module (Cypress, San Jose CA, USA) and converts them to an applied analog voltage via a digital-to-analog converter (DAC). The test chip is connected to the potentiostat via screen-printed electrode (SPE) adaptors attached to the PCB. The SPE adaptors allow for the application of the set voltage to the desired electrochemical system and the recording of the resulting current response, which is then passed back through an analog-to-digital converter (ADC) and transmitted as a user-output signal through the Bluetooth module. The application allows the user to choose between one or two, or three samples and analyzes the electrochemical data accordingly. The user has the option to export the data via email or Messenger on Android and via iCloud as a .txt file on iOS. The Android application was written with Java in Android Studio and the iOS application was written with Swift using SwiftUI in XCode. All electrical components of the potentiostat, including the PCB, are integrated into a custom 3D-printed housing unit designed with a handheld size (101.25 mm x 86.55 mm x 48.99 mm) to allow for true portability and user-friendly manipulation. For more details about the designs and layouts, readers are referred to open-source files accessible through **GitHub link 1**.

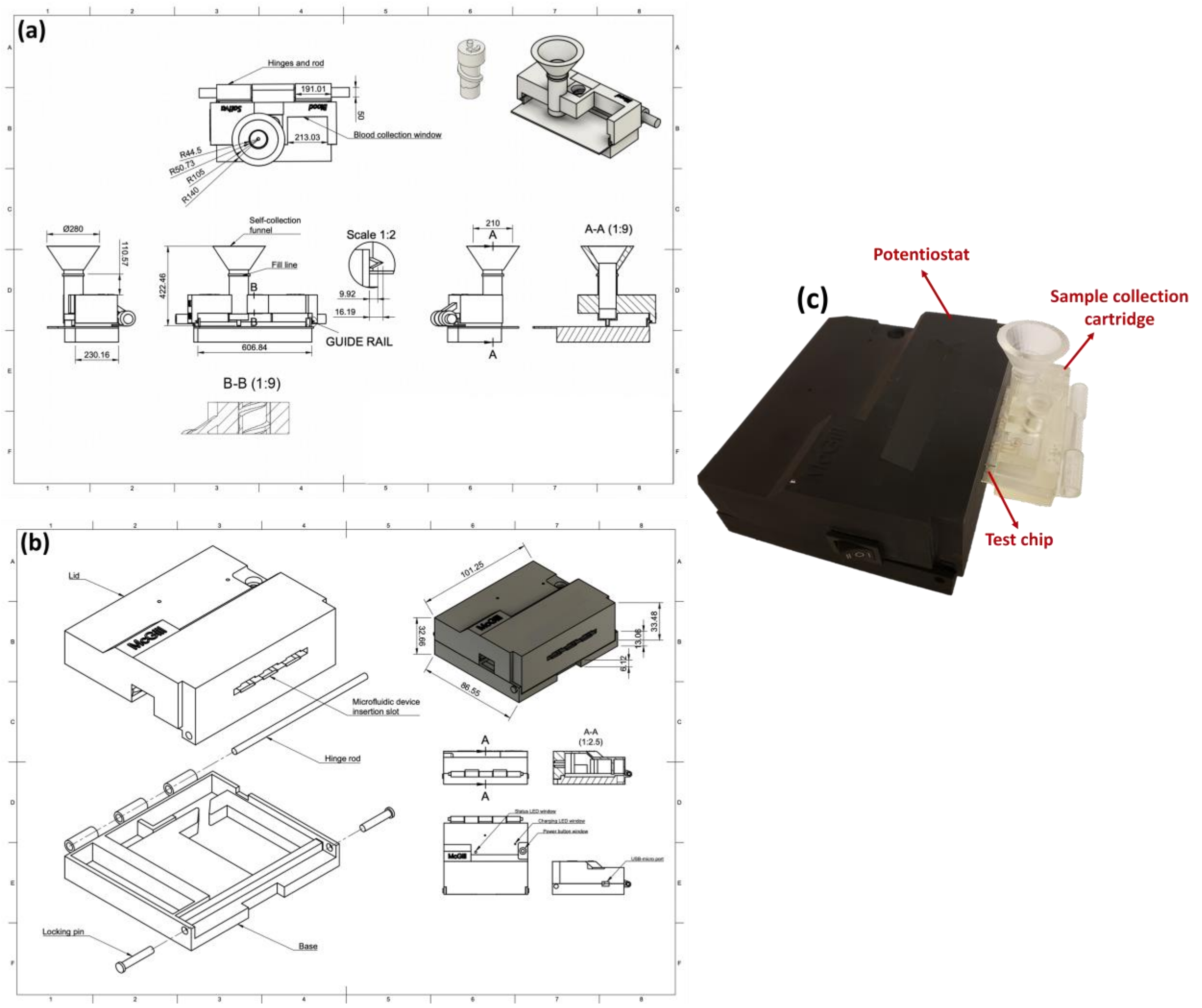


**Fig. S3. Technical drawings of AMMED sample collection cartridge and potentiostat.** a) Technical drawings of AMMED sample collection cartridge, showing all dimensions in 10x millimeters with sectional views representing the hollow regions (clear) and dense regions (shaded), and side views as line drawings. Scale for the drawings is 1:9 unless otherwise indicated, b) Technical drawings of AMMED potentiostat, showing all dimensions in millimeters with sectional views represented by hollow regions (clear) and dense regions (shaded), side views as line drawings, and assemblies as dotted lines. The scale for the drawings are 1:9 unless otherwise indicated, and c) Real image of the AMMED device.

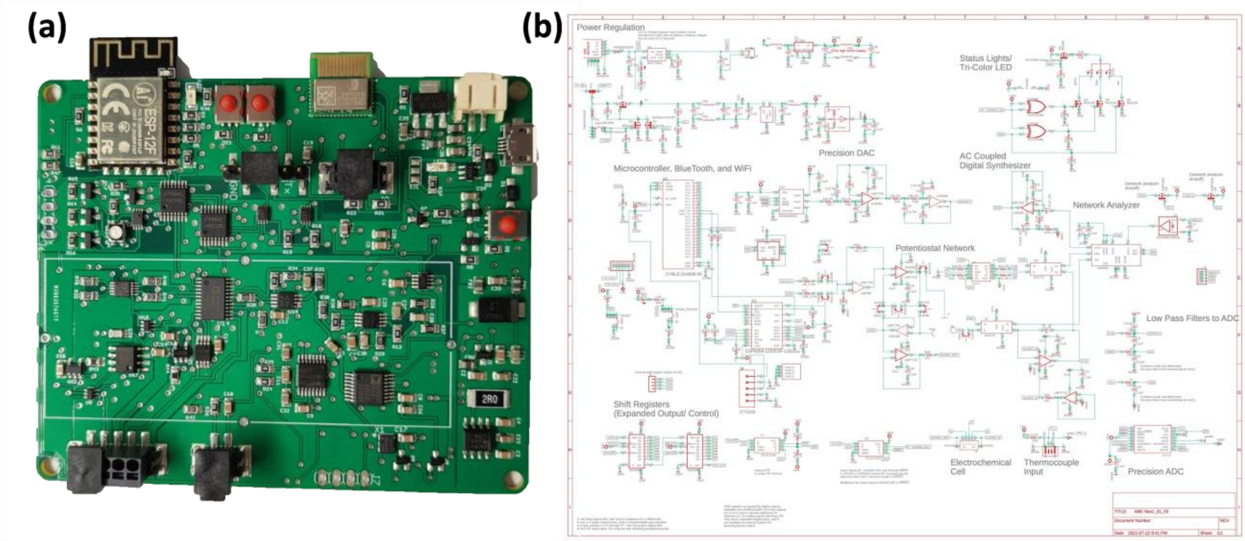


**Fig. S4. Printed circuit board of AMMED potentiostat.** (a) Labeled diagram of the printed circuit board's primary functional components, and (b) Autodesk EAGLE blueprint of the electrical connections of the printed circuit board.

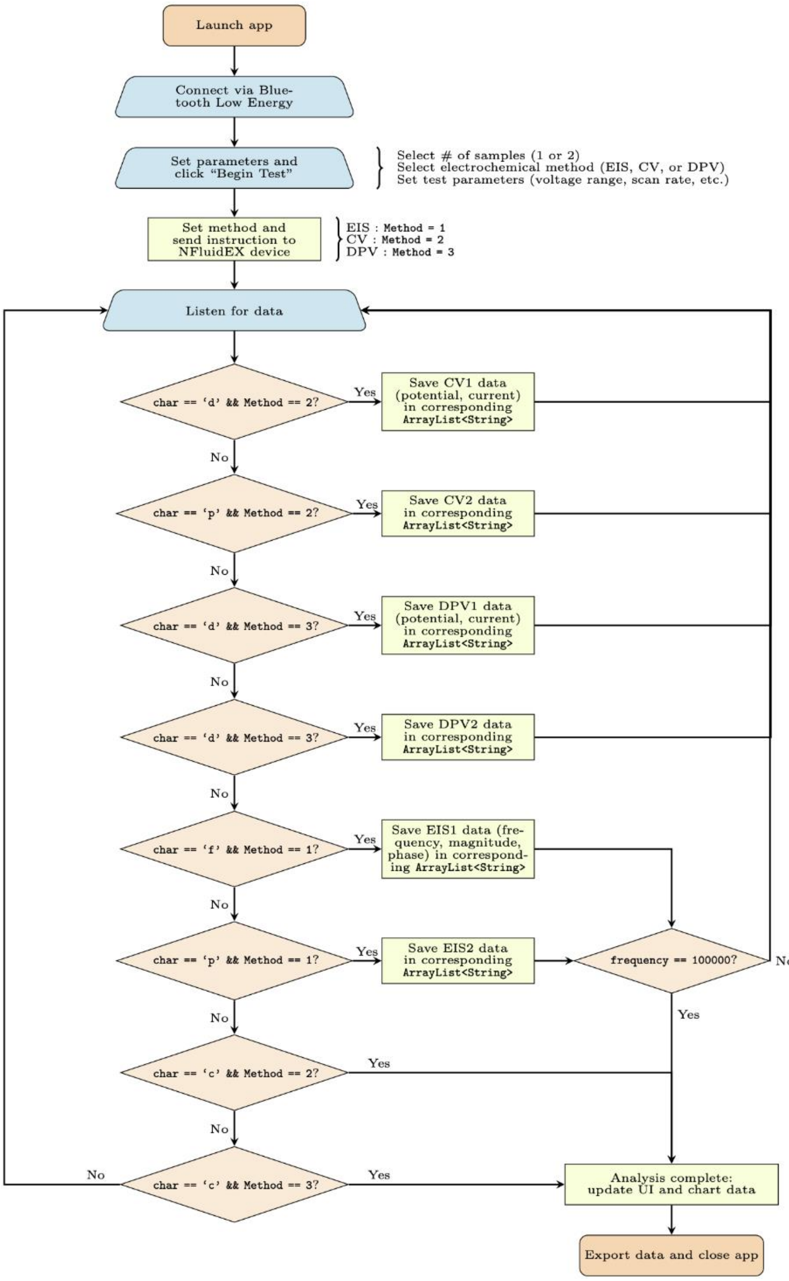


**Fig. S5. The flowchart of the AMMED smartphone application.**

### S4. Electrochemical Measurements

CV measurements were performed at varying scan rates of 12.5, 25, 50, and 100 mV/s (using 5 mM redox probe) and different probe concentrations of 0.5, 1, 2, and 4 mM (at 50 mV/s). DPV measurements were conducted by scanning from -0.2 to +0.6 V with equilibration time 10 s, step potential 2 mV, pulse amplitude 25 mV, pulse width 40 ms, and scan rate 25 mV/s, with varying redox probe concentrations at 2, 4, and 5 mM. EIS was additionally performed using both potentiostats with probe concentrations of 0.5, 1, 2, and 4 mM, with parameters as follows: frequency range 0.1 to 100000 Hz and potential amplitude 100 mV. To obtain charge transfer resistance values ($R_{ct}$), EIS measurements were input to a Python script that fits the data to an equivalent Randles circuit. A combination of EIS and DPV were employed for surface characterization (EIS, DPV) and biosensing (DPV) purposes. Before each electrochemical measurement, the surface of the WEs were washed with PBS and DI water and subsequently dried.

**S5. Fluid flow simulations**

From a sectional view (**Fig. S6**), we can assume that modulating the height of the saliva self-collection funnel shall enable columnated fluid pressure ($P$) (**Equation 1**) that depends on gravity ($g$), the fluid density ($\rho$) and the height of the column ($h$) to push saliva through the funnel fittings and across the filter. The pressure drop required for passage over the filter can be modeled as a porous membrane using Darcy's Law (**Equation 2**) with a known flow rate ($Q$), membrane thickness ($L$), dynamic fluid viscosity ($\mu$), permeability ($\kappa$) and cross-sectional area ($A$).[1] Finally, the pressure drop over the sudden contraction fitting is the pressure required to push fluids through the narrowed funnel that fits into the microfluidic inlet (**Equation 3**) with the initial area ($A1$) and final area ($A2$) that results in a contraction. For a desired flow rate of 100 $\mu L\ s^{-1}$ and known values for saliva viscosity, filter thickness and filter permeability,[2] we found a required pressure drop of ~ 60 Pa over the filter given that the funnel was designed with a 9 mm base diameter. In addition, a sudden contraction fitting (diameter: 2 mm, height: 5 mm) introduced an additional pressure drop that is calculated by considering the mechanical energy lost through friction, and from the previously established flow rate and the given saliva density, the pressure drop over the fitting is ~ 50 Pa.

$$P = \rho g h \tag{1}$$

$$\Delta P = Q\frac{\mu L}{\kappa A} \tag{2}$$

$$\Delta P = \frac{1}{2}\left(0.45\left(1-\frac{A_2}{A_1}\right)\right)\rho\left(\frac{Q}{A_2}\right)^2 + \rho g \Delta h \tag{3}$$

To validate the proposed dimensions, saliva flow through the self-collection funnel was tested through a sequential variation of the fluid height by altering the saliva volume in the funnel. Then, the saliva flowing out of the funnel can be collected over a fixed time interval (e.g. 10 seconds) and the volume can be determined using a graduated pipette to provide an estimate of the acquired volume over time. This has certain assumptions like the expectation of a consistent flow rate over time; we can assume that the flow profile remains constant over the short 10-second interval. When plotted against the simulated values proposed by **Equations 1-3**, we observe a similar linear trend between fluid height and flow rate (**Fig. 3e(ii)**); discrepancies are likely attributed to the higher viscosity of saliva during fresh collection which may have stalled the flow rate.[3] Nonetheless, the linear regime validates the use of columnated pressure height to facilitate the flow of viscous saliva through the self-collection funnel, the filter and the contraction in order to pool at the microfluidic inlet in a passively-driven manner.

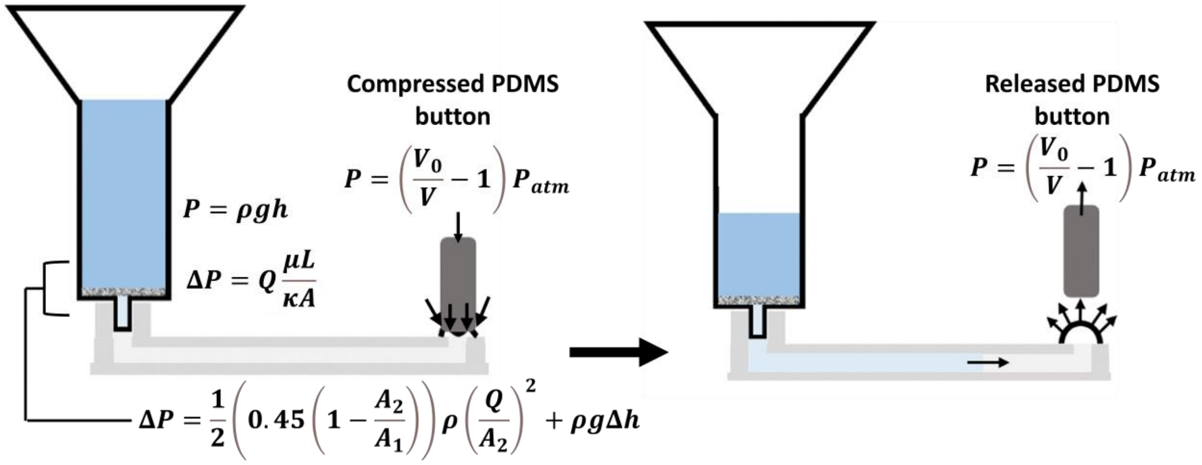


**Fig. S6. Quantitative characterization of the flow rate profile of the passively-driven fluid displacement in the funnel column.**

The flow profile of suction-driven methods can be validated using finite element analysis on COMSOL Multiphysics with geometries imported from AutoCAD. In this simulation, two microchannels are considered: the multiplexed blood channels and the single saliva channel. Suction-based flow relies on the compressed volume of the flexible buttons (**Equation 4**), so the pressure ($P$) depends on the volume of the uncompressed button ($V_0$), the volume of the compressed button ($V$), and the ambient atmospheric pressure of the environment ($P_{atm}$). If we assume that the suction buttons are compressed almost completely (±5% uncompressed), then the pressure drop in the channel is 5000 kPa, which is the input negative pressure at the outlet of the microchannels. Next, we can input the material of the flown biofluids in COMSOL; this includes simulated fluid properties for blood (density = 994 kg $m^{-3}$, dynamic viscosity = 0.004 Pa.s) and saliva (density = 1012 kg.$m^{-3}$, dynamic viscosity = 0.00157 Pa.s). [4-6] The analysis assumed the creeping flow of incompressible fluids with unchanging material properties in a stationary simulation. The resultant velocity distribution in the 3D channels show the xy-plane yielding high-velocity profiles in the channels and lower velocity in the detection chambers (**Fig. 3e(i)**). A surface integration of the velocity profile in the channels yielded a volumetric flow rate of 0.801 µL.s$^{-1}$, which is reasonable given that the channels have volumes of ~1 µL.

$$P = \left(\frac{V_0}{V} - 1\right) P_{atm} \quad (4)$$

Here, $P$ is the negative pressure caused by the suction-button deformation, $V_0$ is the volume of the undeformed pressure chamber, $V$ is the volume of the deformed pressure chamber and $P_{atm}$ is the atmospheric pressure.

This analysis can be extended to view the yz-plane to determine the effect of the fluidic flow on the electrochemical assay in the microchannels. Since the assay is hierarchical, the cross-sectional profile was constructed based on scanning electron microscopy views of the enhanced WE.[7] The assay height and the RE/CE connection was modeled to be 2 µm and 5 µm, respectively; a 50 µm by 25 µm region of interest was studied. From this sectional profile, we observed that the velocity remains at its lowest near the surface of the assay, so flow does not impact the adhesion of the assay or disturb the nano/microstructures (**Fig. 3e(iii)**). The higher velocity at the center of the cross-section demonstrated a parabolic velocity profile, which was indicative of Hagen-Poiseuille flow, so the creeping flow assumptions were valid. The corresponding pressure distributions demonstrated the effect of the RE/CE connection as shielding the assay from high pressure; in the proximity of the RE/CE connection, the incident pressure on the assay was dampened and remained relatively consistent throughout the rest of the cross-sectional plane (**Fig. 3e(iv)**).

**Table S1.** Comparison of existing commercial and hoAMMEDde potentiostats

| Potentiostat | Affordability | Electroanalytical techniques | Multiplexing | Android | iOS | BLE/ Bluetooth | Power source | Ref. |
|---|---|---|---|---|---|---|---|---|
| ABE-Stat (2019) | 105 USD | Voltammetry, EIS | No | Yes | No | Bluetooth | Rechargeable battery | 8 |
| UWED (2018) | 60 USD | Voltammetry | No | No | Yes | BLE | Rechargeable battery | 9 |
| DStat (2015) | 120 CAD | Voltammetry | No | No | No | USB | USB | 10 |
| CheapStat (2011) | 80 USD | Voltammetry | No | No | No | USB | AA batteries or USB | 11 |
| PalmSens EmStat MUX8-R2 | 5200 CAD | LSV, CV, DPV, SWV, NPV, OCP, CA, ZRA, CC, MA, PAD, MPAD | Yes | Yes | No | Bluetooth or USB | USB | PalmSens Inc. |
| Bi-ECDAQ (2022) | 80 CAD | EIS | Yes | No | No | BLE or USB | Rechargeable battery | 12 |
| Enactsense (2022) | 100 USD | Voltammetry | Yes | Yes | No | Bluetooth | N/A | 13 |
| ACEstat (2023) | 60 USD | Voltammetry, EIS | Yes | Yes | No | USB | USB | 14 |
| AMMED (modified ABE-Stat) | 199 CAD | Voltammetry, EIS | Yes | Yes | Yes | BLE | micro-USB | **This work** |

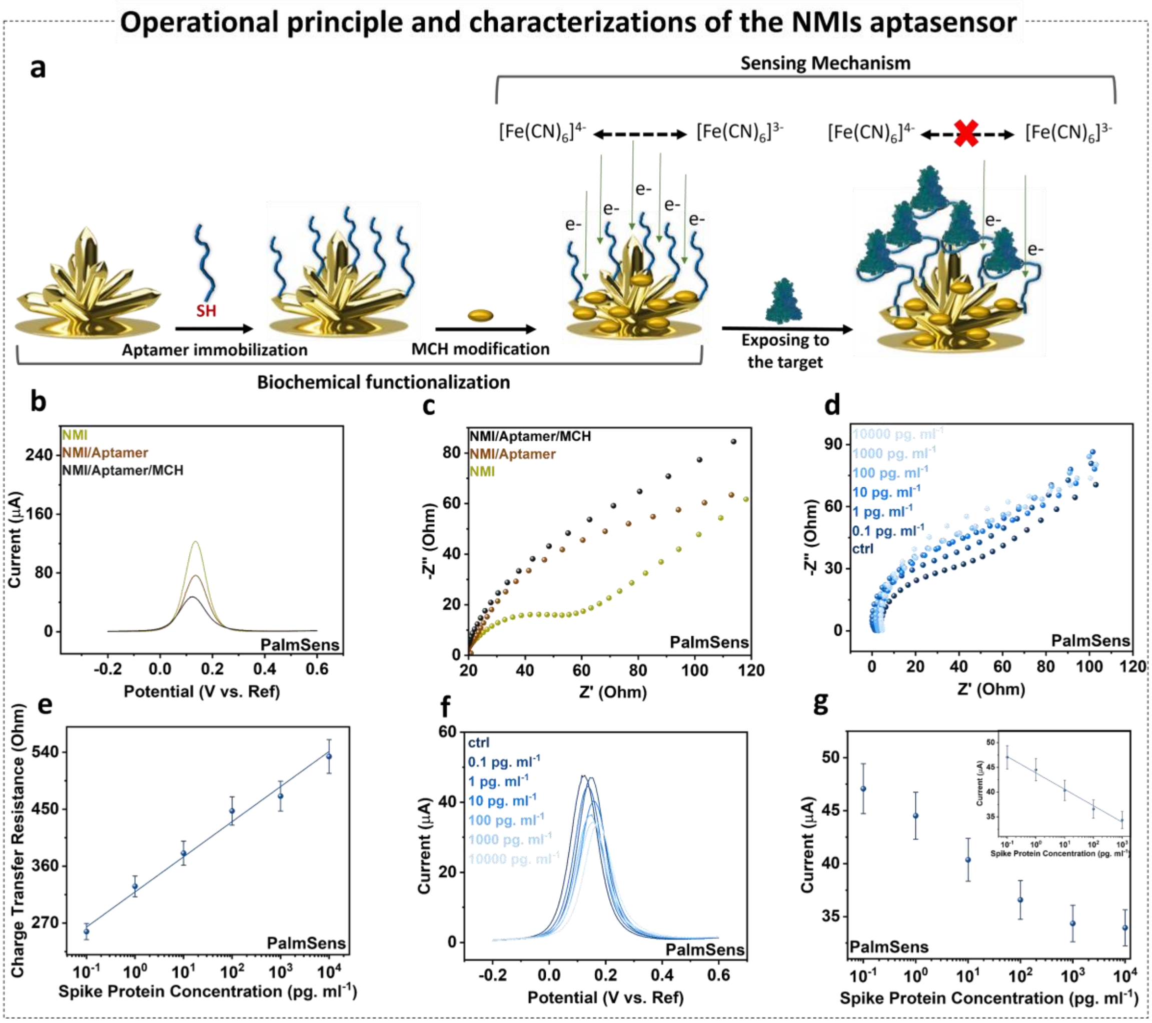


**Fig. S7. Operational principle and characterization of the GNS aptasensor.** (a) Schematic representation of the GNS aptasensor preparation, including the aptamer immobilization and MCH modification, followed by the aptasensor exposure to SARS-CoV-2 S-protein for biosensing evaluation. The (b) DPV, (c) EIS responses associated with each biochemical functionalization step. Electrochemical sensing of SARS-CoV-2 S-protein in PBS spiked with different concentrations of SARS-CoV-2 S-protein via (d) EIS with (e) the corresponding linear calibration plot for the change in charge transfer resistance, and (f) DPV with (g) the corresponding linear calibration plot for the change in peak currents. Data shows mean values ± standard deviation (n=3).

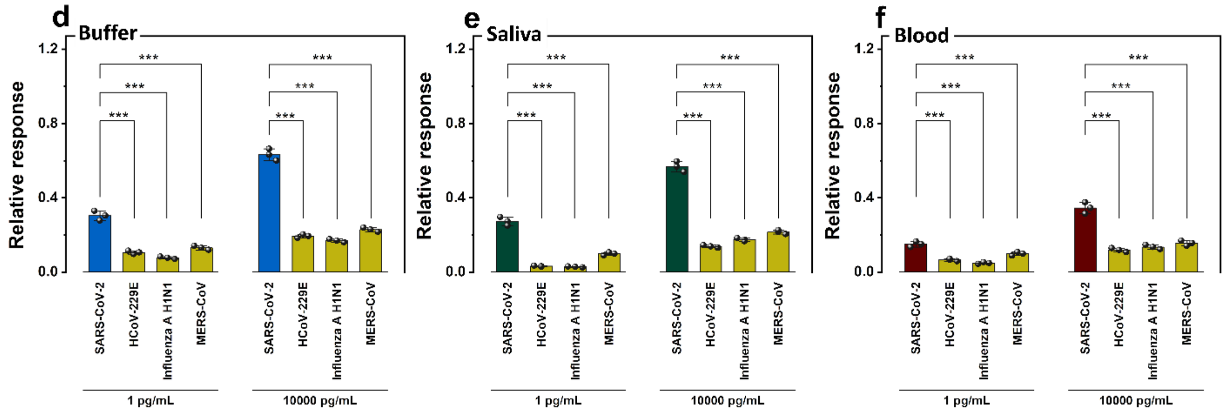


**Fig. S8. Analytical selectivity performance metrics of the GNS aptasensor.** (a) Quantification of the cross-reactivity of the SARS-CoV-2 S-protein as opposed to different viral spike proteins in (a) buffer (b) saliva and (c) blood *** $p < .001$. Data shows mean values ± standard deviation (n=3).

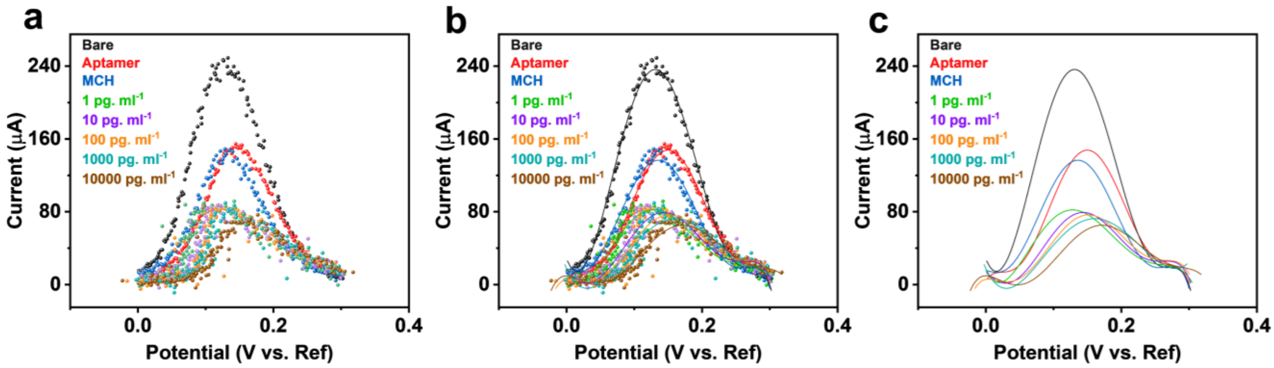


**Fig. S9. Data smoothing approach used for AMMED biosensing.** (a) raw data obtained from AMMED device, (b) 6-degree polynomial curve fitting, (c) fitted 6-degree polynomial trendlines.

### S6. Statistical Analysis

Results were presented as the mean ± the standard deviation over triplicate readings. Statistical analysis was performed using OriginPro (OriginLab, 2021). It is usual that in electrochemical biosensors, the relationship between the concentration (C) and the corresponding current ($I_c$) deviates from linearity on a non-logarithmic scale. Therefore, we need to perform nonlinear regression to obtain the LOD with expression of $3.3sb/a$. In this case, $a$ will be the slope measured at very low concentration (the first two points) and $3.3sb$

will be the 99.5% confidence interval of the *b* parameter. Statistical significance was calculated by performing a one-wayANOVA with post hoc Holm–Sidak's test for mean comparison. The differences between datasets were considered statistically significant for $p < .001$. The figures were generated using the Paired Comparison Plot (version 3.60, OriginLab) graphing application using conservative p values.

**S7. COVID-19 Patient Samples**

10 human saliva samples (5 samples from adult patients with COVID-19 symptoms, such as fever, fatigue, and dry cough, and tested with RT-qPCR) and 5 samples from healthy controls were supplied by the University Health Network's PRESERVE-Pandemic Response Biobank for testing on the assay (REB #20-5364). Free authorization and consent forms were signed by patients, and their clinical samples were collected according to the laboratory regulation. The samples were assessed at a Level 2+ facility situated in the Montreal Jewish General Hospital.

a cycle threshold (Ct) value for RNA-dependent RNA polymerase (RdRp) amplification in RT-qPCR was used to determine the correlation between SARS-CoV-2 viral particle concentrations and RT-qPCR responses in order to quantify the AMMED results in comparison to the gold standard RT-qPCR method. As illustrated in **Fig. S10**, the calibration plot relates RT-qPCR Ct values to viral particle concentration, which is consistent with the known trend of a decreasing Ct with increasing viral load. [15, 16]

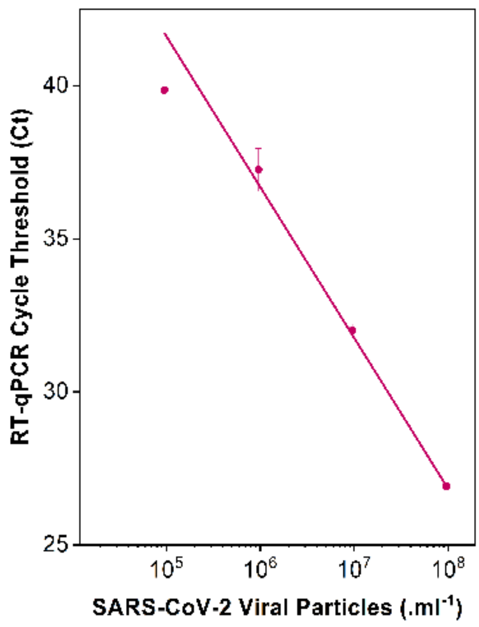


**Fig. S10.** Calibration plot of RT-qPCR Ct values as a function of heat-inactivated SARS-CoV-2 viral particle concentration.

Viral particle concentration refers to the number of intact and infectious virus particles in a sample, varying based on infection stage, replication rate, and immune response. The relationship with spike protein levels is not always direct; increased viral particle concentration may not always proportionally correlate with spike protein levels. Factors like viral replication efficiency, spike protein shedding, and immune responses can influence spike protein levels independently of viral particle concentration. AMMED demonstrated a significant correlation with conventional RT-qPCR analysis, revealing varying Ct values from 22.48 (Patient 5) to 29.28 (Patient 3). Remarkably, AMMED exhibited lower currents for patient samples with lower Ct values. This indicates a potential association between higher viral loads and increased electrochemical signal response. These findings underscore the potential of AMMED as a sensitive and complementary method for viral load quantification and detection in clinical samples (**Table S2**).

**Table S2**. RT-qPCR Ct values for patient samples

| Patient Code | Ct value |
|---|---|
| Patient 1 | 27.58 |
| Patient 2 | 28.13 |
| Patient 3 | 29.28 |
| Patient 4 | 22.88 |
| Patient 5 | 22.48 |

**Table S3.** Cost breakdown for AMMED device

| Part | Bulk cost | Quantity | Cost per unit | Vendor |
|---|---|---|---|---|
| **AMMED Test chip** | | | | |
| Microfluidic device fabrication | 3D printing of microfluidic part | ~ 1 hours total | $0.5 | FormLabs 3D printer |
| Aptamer-based assay | $250 per 100nmole | 3 electrodes per device | $0.0405 | Integrated DNA Technologies IDT |
| Suction buttons | $109 per 0.5 kilograms | 1 mL per device | $0.21 | Dow SYLGARD |
| **AMMED Sample collection cartridge** | | | | |

| | | | | |
|---|---|---|---|---|
| 3D printing | $24.95 per 800 mL | 16 mL per device | $0.50 | Filaments.ca, Formlabs |
| Whatman Grade 4 filter | $1.15 per 10 filters | 1/4 filter per device | $0.029 | Whatman |
| **Total** | **$1.03 per AMMED sample collection cartridge and test chip** | | | |
| **AMMED potentiostat** | | | | |
| 3D printing | $24.95 per 800 mL | 240 mL per device | $7.49 | Filaments.ca, Formlabs |
| Printed circuit board (fully assembled) | $92.85 per board | 1 per device | $92.85 | PCBWay |
| Battery | $14.99 per battery | 1 per device | $14.99 | Canada Robotix |
| Relay module | $26.49 per 5 modules | 1 per device | $5.30 | Amazon |
| Screen-printed electrode adaptors | $20 per adaptor | 3 per device | $60 | IORodeo |
| 3-way manual toggle switch | $6.15 per 5 switches | 1 per device | $1.23 | Amazon |
| 3-way servo splitter cable | $16.99 per cable | 1 per device | $16.99 | Amazon |
| Jumper wires | $11.98 per 240 wires | 3 per device | $0.15 | Amazon |

| Total | $199 per AMMED potentiostat |
|---|---|

**S8. GitHub open-source links**

Link 1: GitHub - Potentiostat/AMMED

Link 2: GitHub - iOS/AMMED

Link 3: GitHub - Android /AMMED
